# CryptoBench: Cryptic protein-ligand binding sites dataset and benchmark

**DOI:** 10.1101/2024.08.20.608828

**Authors:** Vít Škrhák, Marian Novotný, Christos P. Feidakis, Radoslav Krivák, David Hoksza

## Abstract

Structure-based methods for detecting protein-ligand binding sites play a crucial role in various domains, from fundamental research to biomedical applications. However, current prediction methodologies often rely on holo (ligand-bound) protein conformations for training and evaluation, overlooking the significance of the apo (ligand-free) states. This oversight is particularly problematic in the case of cryptic binding sites (CBSs) where holo-based assessment yields unrealistic performance expectations. To advance the development in this domain, we introduce CryptoBench, a benchmark dataset tailored for training and evaluating novel CBS prediction methodologies. CryptoBench is constructed upon a large collection of apo-holo protein pairs, grouped by UniProtID, clustered by sequence identity, and filtered to contain only structures with substantial structural change in the binding site. CryptoBench comprises 1,107 structures with predefined cross-validation splits, making it the most extensive CBS dataset to date. To establish a performance baseline, we measured the predictive power of sequence- and structure-based CBS residue prediction methods using the benchmark. We selected PocketMiner as the state-of-the-art representative of the structure-based methods for CBS detection, and P2Rank, a widely-used structure-based method for general binding site prediction that is not specifically tailored for cryptic sites. For sequence-based approaches, we trained a neural network to classify binding residues using protein language model embeddings. Our sequence-based approach outperformed PocketMiner and P2Rank across key metrics, including AUC, AUPRC, MCC, and F1 scores. These results provide baseline benchmark results for future CBS and potentially also non-CBS prediction endeavors, leveraging CryptoBench as the foundational platform for further advancements in the field.

## Introduction

Proteins serve as the molecular workhorses of living organisms, executing a myriad of functions essential for life. Their three-dimensional structure, which serves as the blueprint for their biochemical activities, is central to the functionality of proteins. Protein structure embodies a dynamic landscape characterized by conformational flexibility and adaptability in response to environmental triggers. In this context, ligand binding sites, regions where a protein binds to its interacting partners, deserve special attention. Specifically, cryptic binding sites (CBSs) are ligand binding sites that are not readily apparent or accessible in their ligand-free (apo) state but become exposed due to external triggers, thus enabling the binding of a ligand and forming the ligand-bound (holo) state. Therefore, CBSs can be loosely defined as sites identifiable in the ligand-bound but not in the unbound structure [1].

Due to their importance as potential targets for drug discovery and protein engineering, binding sites, in general, have gathered considerable attention, resulting in the development of a wide range of binding site prediction methods [2]. As the structure primarily drives the interaction, the structure-based approaches are typically considered superior to sequence-based methods. However, structure-based approaches are fallible with respect to protein conformational flexibility as they are developed/trained to recognize certain structural characteristics. For instance, the authors of P2Rank [3] quantified the effect of different features on the prediction quality, showing that the biggest effect by far comes from the protrusion of surface regions. The reliance on structural properties becomes problematic in the case of CBSs as their shape can differ significantly between the apo and holo forms, resulting in decreased performance when measured on the apo form of the proteins [4, 5]. This observation led to the development of methods specialized in predicting CBSs. These methods are based on molecular dynamics [6, 7, 8, 9, 10], machine learning (ML) [11, 5] or a combination of both [12]. A commonly used dataset to evaluate CBS prediction approaches is the one introduced with the CryptoSite method [13]. The CryptoSite dataset, however, faces several shortcomings: i) size, ii) definition of crypticity, and iii) lack of pockets with substantial conformational changes. Regarding size, the CryptoSite dataset comprises only 93 cryptic binding pockets. As for the second point, the CryptoSite dataset construction methodology utilizes two other protein-ligand prediction tools, FPocket and ConCavity, and declares a pocket cryptic when detected in its holo but not in its apo form by these tools. A concern with such an approach is that biases within these predictors can leak into the dataset and, therefore, propagate if the dataset is further employed for testing or training new software. Finally, the CryptoSite dataset includes a relatively small number of cryptic pockets requiring large conformational changes [11], leading to a potential underrepresentation of such pockets within the dataset. The recently introduced PocketMiner dataset [11] does not suffer from the possibility of introducing biases and also comprises apo-holo pairs with large structural rearrangements but comes with a significant trade-off in size, containing only 39 cryptic pockets.

As shown above, the current datasets are quite small, which is understandable considering that building a dataset on a large scale is a complex endeavor. It requires identifying apo-holo pairs for several ligand binding sites across the entire PDB, classifying them as cryptic or non-cryptic (i.e., by using a suitable metric), and selecting a representative apo candidate in the presence of multiple. However, without such a dataset, a substantial volume of cryptic pockets within our observed proteome remains unexplored and underrepresented. To address these issues, we present CryptoBench, the most comprehensive dataset of cryptic binding sites so far. In this work, we understand a binding site to be cryptic when there is a significant structural change between the apo and holo form (For motivation and definition, see the Dataset Construction section, and for further discussion, refer to the Discussion section). By introducing CryptoBench, we aim to expand the coverage of cryptic binding sites from protein structures available in the PDB [14] thereby facilitating the detection of cryptic binding sites, a promising strategy for drug repurposing [15].

### Dataset Construction

The assembled dataset of cryptic binding sites is built on top of AHoJ-DB [16], a database of precomputed apo-holo results of the AHoJ tool [17]. AHoJ-DB links apo and holo states of a ligand binding site, which are derived from different structures in the PDB. AHoJ defines the binding site by considering protein residues within a user-defined distance threshold (default 4.5 Å) from the atoms of the specified ligand. At the time of writing (August 2024), AHoJ-DB featured apo-holo and holo-holo pairs for 522,153 biologically relevant protein-ligand interactions [18] across the entire PDB. AHoJ-DB is constructed by mapping the binding residues of each specified protein-ligand interaction across all other structures in the PDB that are assigned the same UniProt accession, and registering these candidate pockets as holo or apo depending on the presence or absence of bound ligands. Apo and holo structures are mapped residue-wise by linking the common binding residues by their UniProt sequence indices, thereby tracking the same pocket across multiple structures. Several metrics are reported for each pocket (including RMSD, volume, solvent accessible surface area (SASA), molecular surface) and the chains that comprise them. The resource features a substantial number of both holo-holo (∼43 million) and apo-holo pocket pairs (∼14 million) that are, however, not explicitly annotated as cryptic or non-cryptic. Furthermore, of the 522,153 holo pockets cataloged in AHoJ-DB, approximately 53,6% (280,058 holo pockets) lack a corresponding apo structure. While some of these pockets may potentially be cryptic, without sufficient data for comparison of the apo and holo form, we cannot classify them as such.

Although, for a single protein-ligand structure, AHoJ-DB includes all apo and holo structures, when assembling the dataset, we considered only the apo states, i.e., disregarding the alternative holo structures. Subsequently, we filtered the apo-holo pairs down by keeping only pockets from structures with a resolution of 2.5 Å or higher. Further, we refined the results by only keeping apo pockets where the number of binding residues is equal in both apo and holo states. After conducting these steps, the filtered subset of AHoJ-DB totaled 4,683,968 pairs, each representing a single binding pocket in its apo and holo state.

#### Selecting a suitable crypticity metric

To differentiate cryptic and non-cryptic apo-holo pairs, a suitable metric to distinguish between regular and cryptic binding sites has to be selected. Such a metric can then be utilized to extract a dataset comprising solely cryptic pockets from the broader AHoJ-DB dataset. The primary metrics we considered included pocket solvent accessible surface area (SASA) [19], pocket molecular surface [20], pocket volume [21], and all-atom pocket RMSD (see supplementary information for the difference between SASA and molecular surface). All the metrics were evaluated exclusively within the scope of the pocket, i.e., the metrics were computed using only the pocket residues. These metrics were chosen for their potential to capture the dynamic nature of cryptic pockets, as opposed to the static nature of regular pockets within the apo-holo pairs. The metric values for each apo-holo pair were extracted from AHoJ-DB.

To identify the most appropriate metric and its threshold for dataset filtering, we conducted a comparative analysis between two datasets: one extracted from AHoJ-DB^1^, representing a general dataset containing a mixture of regular and potentially cryptic binding sites, and the other being the PocketMiner test dataset [11], which contained 38 hand-picked structures with exclusively cryptic binding sites. While the metrics values for the records from AHoJ-DB were already precomputed, we additionally computed the metrics values for the records from the PocketMiner test dataset. Subsequently, the metric values for every pocket in both datasets were analyzed to assess their suitability for distinguishing between regular and cryptic binding sites. The resulting violin plots in Figure 1 illustrate the behavior of each metric within the context of these datasets. It is important to note that while molecular surface, SASA, and volume yield two values for each pair — one for apo and the other for holo — the RMSD yields only a single value for each pair. Therefore, in the context of the violin plots, while RMSD values were used as they were, the molecular surface, SASA, and volume values were computed as the difference between the apo and holo values, normalized over the number of pocket residues (pocket length):

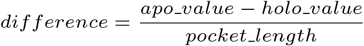

**Fig. 1.**
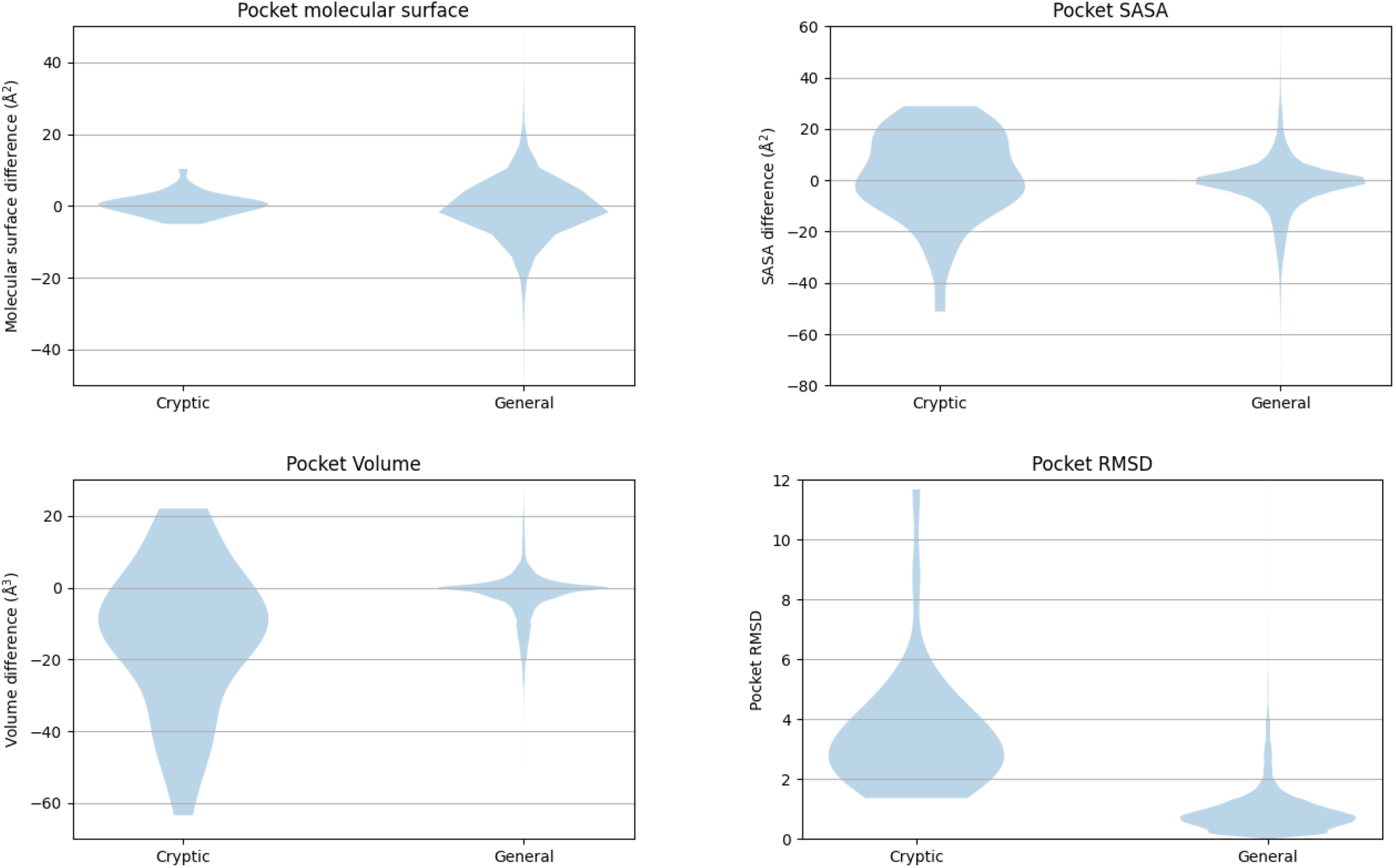
The distribution of four metrics — pocket SASA, pocket molecular surface, pocket volume, and pocket RMSD — across two datasets: one representing cryptic-only binding sites (PocketMiner test dataset) and the other representing general binding sites (AHoJ-DB dataset). Overall, the violin plots for pocket molecular surface largely overlap, indicating similar distributions between the two datasets. In contrast, the violin plots for pocket volume and pocket SASA show partial overlap, suggesting some differences in distribution between cryptic and general binding sites. Notably, the violin plots for pocket RMSD exhibit nearly no overlap, with a clear borderline between 1.5 and 2 Å.

As observed from the violin plots, the molecular surface exhibits significant overlap in its distributions, suggesting it is not practical for distinguishing between cryptic and regular binding sites. While there is less overlap in the distributions of pocket SASA and pocket volume, a clear boundary between the values of cryptic and regular datasets is not apparent. Finally, pocket RMSD shows promising results, with values for the general dataset typically not exceeding 1.5 Å, and values for the cryptic dataset generally remaining above 2 Å.

Thus, out of the considered metrics, pocket RMSD seems to be the most effective metric for distinguishing between cryptic and regular binding sites. Given the insights provided by the violin plots in Figure 1, we established pocket RMSD bigger than 2 Å as a suitable threshold for differentiating between cryptic and regular binding sites. By selecting the upper bound of the borderline interval [1.5, 2] Å, we aimed to minimize the risk of unintentionally including regular binding sites in the cryptic dataset, particularly when filtering a general dataset to create a cryptic one.

To further evaluate the selected threshold, we looked at the RMSD values within the PocketMiner dataset of cryptic binding sites. We found that 32 out of 38 pairs have higher than 2Å RMSD and are, therefore, in agreement with our threshold. Furthermore, we manually inspected the remaining six pairs with RMSD between 1,35 and 1,93Å. There were four structure pairs with larger rearrangements of the binding sites - see, for example, the surface representation of the 4i92 (apo) and 4i94 (holo) structure pair in Figure 2. The substantial part of the binding sites is not present in the apo form due to the conformational change of side chains of Asn71 and Arg186. There are, however, also two structure pairs where the conformational change is less substantial (4w51-4w58, 3ppn-3ppr) - see supplementary data for examples. However, the degree to which these binding sites should be classified as cryptic remained uncertain. Ultimately, these findings indicated that although our selected threshold can miss some cryptic binding sites, our decision to set the threshold at 2 Å is reasonable.

**Fig. 2.**
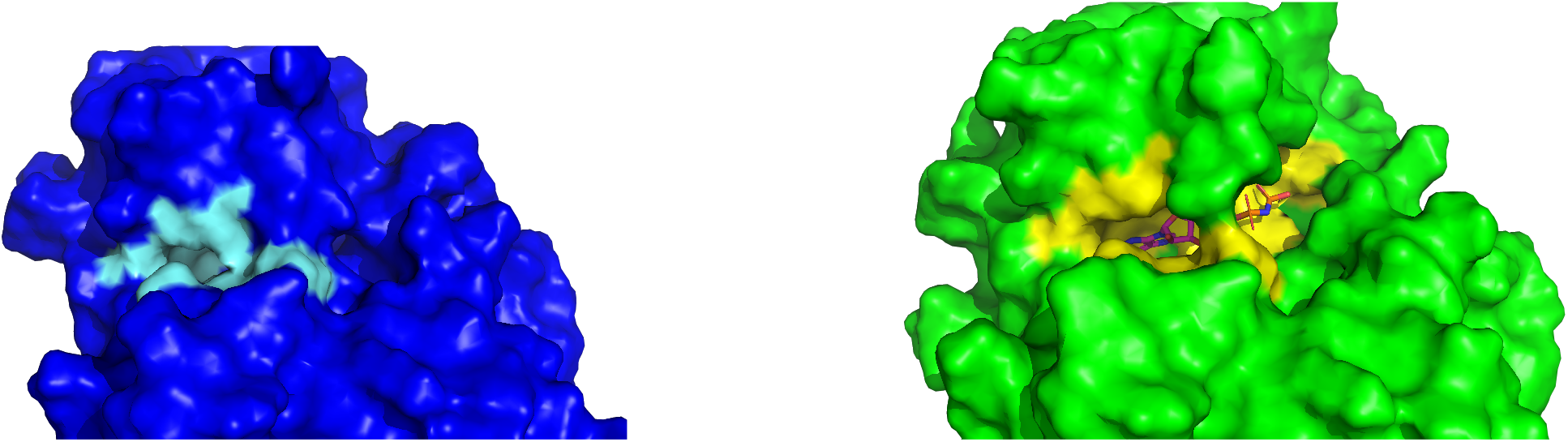
A comparison of the apo and holo cryptic binding site. A) The holo structure of plant pseudokinase binding AMP (4i94) in surface representation. B) The apo structure of the same protein (4i92) in surface representation – the right part of the binding site is altered.

From the previous discussion follows the definition of a cryptic binding site for the purpose of the CryptoBench dataset. The cryptic binding site refers to a region in a protein that can bind a ligand and undergoes a significant structural change between its holo (ligand-bound) and apo (unbound) forms. A significant change is defined as a difference of at least 2 Å RMSD between the binding residues in the apo and holo forms. Binding residues are defined as those located within a 4.5 Å radius from the ligand in the holo form. The corresponding binding residues in the apo form are identified by mapping the holo-form binding residues onto the apo structure using UniProt mapping.

#### Apo-holo pairs pool filtering

Considering the lack of a consensus definition for binding site crypticity, and the inclusivity of AHoJ-DB in terms of conformational changes that occur within a given protein (UniProt ID), it is important to establish not only a minimum threshold for detecting significant conformational changes in the binding site (i.e., crypticity), but also an upper limit to ensure that i) both the global structure of the protein, beyond the binding site, remains relatively stable and recognizable between the two states, and ii) the relative position of the binding site does not deviate significantly between the two states. While an argument can be made for the inclusion of any structure of the same UniProt, regardless of the extent of its global conformational changes that can extend to disorder-to-order transitions or domain swaps and hinge-like motions, often the difference between the global similarity (i.e., TM-score) of structures of the same sequence, can be larger than that of different sequences, or of the minimum recommended TM-score of 0.5 for structures that “assume generally the same fold in SCOP/CATH” [22]. Such differences could also prove problematic during visual inspection where there is no visible resemblance between the global structures of the apo and holo states, but perhaps more importantly, the scope of the cryptic binding site can be overstretched or lost when the structural changes span the entire protein chain. In addition, we choose to exclude cases where the compactness of the binding site changes significantly on account of a disordered or unfolded state compared to the typically well-defined and compact holo state. To construct the cryptic dataset, the original 4,683,968 apo-holo pairs retrieved from AHoJ-DB were processed as follows:

1. *resolution filtering:* all records with a resolution worse than 2.5 Å were filtered out; on top of that, all records where pocket length in the apo state and holo state did not match were filtered out,
2. *geometric quality assurance filtering* 2. to ensure that the conformational changes are restricted to the binding site, we establish a threshold for the minimum accepted global similarity between the two states of the protein chain that comprises the binding site, achieved by a minimum TM-score of 0.5. Also, to filter out domain swaps and large intrachain motions, we establish a maximum distance threshold of 4 Å between the centers of apo and holo binding sites, after the global structural alignment. Furthermore, we establish a threshold for the allowed change in the level of compactness of the binding site between the apo and holo states, by allowing up to a 20% change in the radius of gyration from the holo state. Lastly, at least 50 observed residues from the protein must overlap with its UniProt sequence. All metrics were taken from AHoJ-DB,
3. *pocket RMSD filtering* records with pocket RMSD below 2 Å were filtered out,
4. *ligand filtering:* similarly to P2Rank [3], we excluded ligands where the number of atoms is less than 5. Furthermore, the name of the PDB group is not on the list of ignored groups: (HOH, DOD, WAT, UNK, ABA, MPD, GOL, SO4, PO4)^2^,
5. *clustering*: unique UniProt sequences were clustered based on 40% similarity [24],
6. *selection of representatives*: from each cluster from the previous step, we selected one representative apo-holo pair based on maximal pocket RMSD, aiming to include pairs with the most significant structural changes,
7. *searching for additional pockets*: considering that one apo structure can contain multiple cryptic pockets or a single cryptic pocket may bind multiple types of ligands, we conducted a second search within the output of the *pocket RMSD filtering* phase to identify these pockets and include them into the dataset. Therefore, within our dataset, one apo structure can be paired with more than one holo structures.

The number of apo-holo pairs outputted by each filtering phase summarizes Table 1.

**Table 1.** The number of apo-holo pairs left after each filtering phase.

|  | CryptoBench dataset |
| --- | --- |
| *resolution filtering* | 4,683,968 |
| *geometric quality assurance filtering* | 4,423,220 |
| *pocket RMSD filtering* | 221,026 |
| *ligand filtering* | 171,876 |
| *clustering & selection of representatives* | 1,107 |
| *additional pockets search* | 5,493 |

### Dataset Structure

Practically, the resulting dataset comprises a list of apo structures identified by their apo PDB ID, with each PDB ID associated with one or more cryptic pockets. Each cryptic pocket represents a single record in the dataset. Within the context of a single apo structure, each pocket may correspond to a different holo structure and a different holo chain as well. Consequently, each pocket in the apo structure is characterized by the PDB identifier of the holo structure, chain identifier, ligand name, ligand residue number (to distinguish between different ligands of the same type), and both apo and holo pocket residue selections.

### Statistics

The apo-holo pairs may contain multiple cryptic binding sites, each with its unique set of residues. For the sake of statistics, to differentiate between distinct binding sites within a single apo structure, we applied a criterion where two binding sites are considered separate if their residues have less than a 75% overlap. Similarly, we identified cryptic binding sites capable of binding more than one type of ligand - promiscuous pockets. Using the same 75% threshold, we classified a pocket as promiscuous if it overlapped by more than 75% with another pocket binding different ligands. Lastly, we counted the number of cryptic binding sites spanning more than one chain. The statistics are shown in Table 2.

**Table 2.** Statistics for the CryptoBench dataset.

|  | CryptoBench dataset |
| --- | --- |
| *apo-holo pairs* | 1,107 |
| *cryptic pockets* | 1,361 |
| *promiscuous pockets* | 371 |
| *multi-chain pockets* | 197 |
| *avg. pocket size* | $16.60 \pm 7.22$ |
| *avg. # of observed residues* | $290.34 \pm 154.00^3$ |

### CryptoBench benchmark

To be able to use the dataset for fair validation of ML-based prediction approaches, we pre-defined the train-test splits of the CryptoBench dataset. As shown in another study, the methods’ performance and superiority might be influenced by how K-fold splits are defined [25]. Therefore, we also established the K-fold splits for the train set. Furthermore, unlike datasets with other media, such as images, protein datasets are vulnerable to information leakage between splits if not carefully managed [26]. To mitigate the risk of including homologous proteins in different splits, causing information leakage, we conducted another round of clustering on the dataset. This round of clustering utilized a threshold of 10% sequence identity to ensure minimal relation between records from each data split [24]. These clusters were then joined into the splits forming the resulting CryptoBench benchmark.

We used an 80:20 ratio for the train-test split. The resulting train split was further divided into a 4-fold split. The number of structures in each fold and the train/test split is shown in Table 3.

**Table 3.** Number of structures in train/test split. The train set is broken up into folds.

|  | CryptoBench dataset |
| --- | --- |
| TRAIN subset | 885 |
| TEST subset | 222 |
| TRAIN-FOLD-1 subset | 219 |
| TRAIN-FOLD-2 subset | 222 |
| TRAIN-FOLD-3 subset | 222 |
| TRAIN-FOLD-4 subset | 222 |

#### Baseline Evaluation

To validate the dataset’s usability, we used it to evaluate one sequence-based and one structure-based method for detecting cryptic binding residues. We used PocketMiner [11] as the representative of structure-based methods, which, according to the presented experiments [11], has demonstrated superior performance over the previous state-of-the-art tool, CryptoSite [12]. For the sequence baseline, we implemented our own neural network architecture utilizing a protein language model (**pLM-NN**), as a similar architecture proved promising when evaluated on the CryptoSite dataset [5]. Finally, we included P2Rank in the comparison as a representative of non-CBS-specific methods, which has been shown to perform quite well for CBS detection [4, 5].

It should be emphasized that PocketMiner, P2Rank and the pLM-NN models were trained on different datasets. While the pLM-NN model was trained on CryptoBench, we used the model available on the project GitHub page for PocketMiner without any retraining or fine-tuning. Similarly, we utilized P2Rank’s off-the-shelf pretrained prediction model, which was trained on P2Rank’s own training dataset, without any further adjustment. This might skew the performance of PocketMiner and P2Rank as more data might be available for training. On the other hand, we did not control for possible data leakage between PocketMiner/P2Rank training and CryptoBench test sets.

### Protein language model classifier implementation

The implemented method is purely sequence-based, as the input for the pLM-NN consisted solely of protein embeddings from the ESM2-3B model [27, 28], generated from whole UniProt sequences. The ESM2-3B model generates an embedding of size 2560 for each sequence residue. Although the embeddings were computed from the whole sequence, only those embeddings corresponding to the residues observed in the structure were kept for training and evaluation.

The residue labels, indicating whether a particular residue is part of a cryptic site, were generated by merging all available cryptic pockets. Therefore, if a protein structure contains multiple pockets, all of them are incorporated into the labeling, resulting in a binary classification problem where each residue is either binding or non-binding.

Figure 3 illustrates the classification process. First, embeddings were acquired from the ESM2-3B model using the entire UniProt sequence. Subsequently, only the embeddings corresponding to observed residues were kept, while unobserved residues or those not present in the PDB structure were discarded. These filtered embeddings served as input for the pLM-NN. The network outputs the probability of a residue being part of a cryptic binding site. Therefore, the pLM-NN yields results only for the observed residues, with the unobserved residues colored grey in the diagram.

**Fig. 3.**
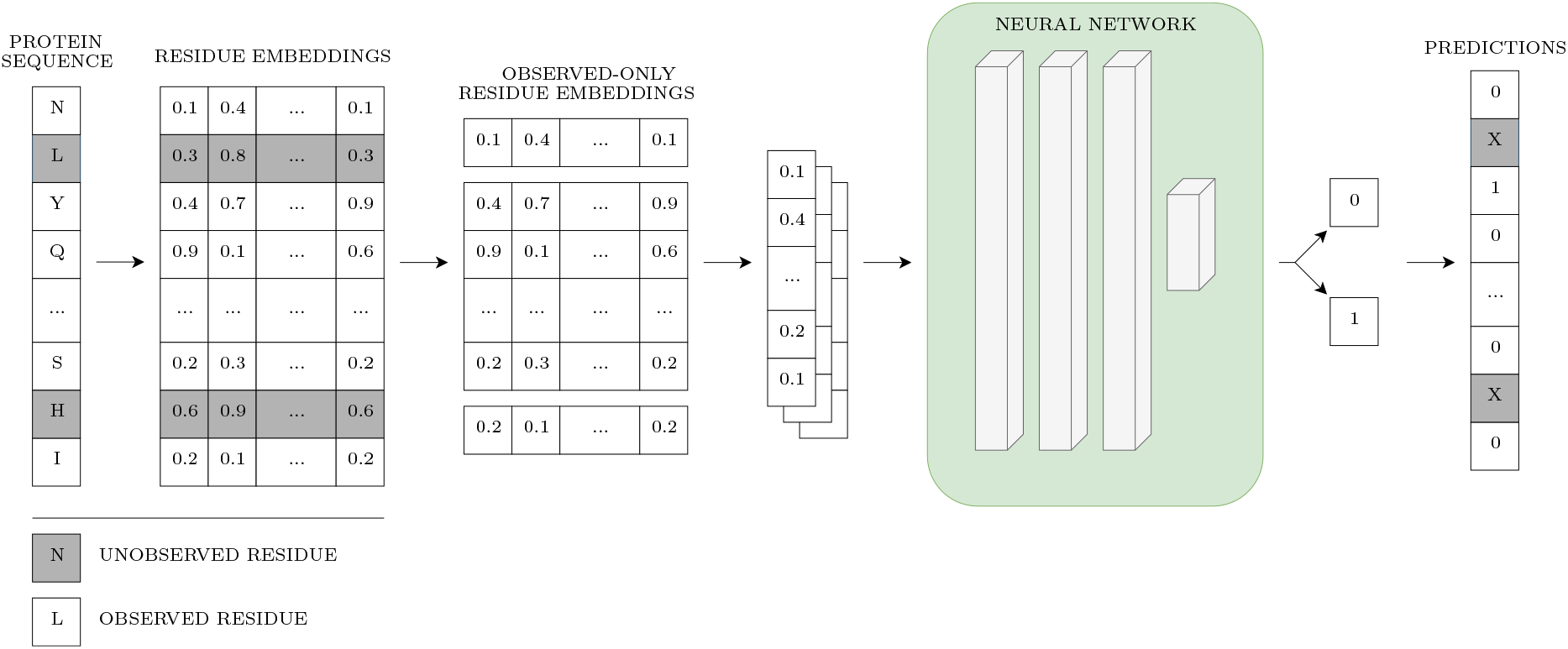
Overview of the pLM-NN. Grey coloring depicts unobserved residues. The prediction was not made for the unobserved residues, as the embeddings for unobserved residues were discarded for the sake of fair comparison with structure-based methods.

To determine the optimal architecture of the pLM-NN, SklearnTuner [29] was used to perform the cross-validated hyperparameter search using the predefined 4-fold train split of the training subset. The final architecture employed an L2 regularizer and ReLU activation. It was composed of 3 layers, each followed by a dropout layer with the dropout rate equal to 0.3. The first two layers each contained 256 neurons, while the last layer consisted of 2 neurons. Overall, the setup yielded 721,922 parameters. The binary cross-entropy loss function was used and optimized using the Adam optimizer, with a learning rate set to 1e-04. The training was executed for 7 epochs, with a batch size 2,048.

## Results

It should be emphasized that the pLM-NN utilized solely sequence information and training was conducted exclusively using observed residues. This was done to ensure a fair comparison between the trained pLM-NN and PocketMiner, which is a structure-based method, therefore it cannot evaluate unobserved residues. Both methods output a probability of a residue being cryptic. We used a 0.75 threshold to assign the residue to either the positive or negative class. As the dataset is heavily imbalanced, a decision threshold of 0.75 favors the negative class, which is significantly larger. In the case of P2Rank, apart from providing the probability of a residue being part of a binding site, the tool also outputs a binary classification of the residue into either positive or negative classes directly, thus no additional thresholding was needed.

Using the aforementioned binary classification, true positive rate (**TPR**), false positive rate (**FPR**), F1 score (**F1**), accuracy (**ACC**), and Matthew’s correlation coefficient (**MCC**) were calculated for each prediction method. In the case of the area under the curve (**AUC**), the probabilities were utilized directly to construct the receiver operating characteristics curve from which AUC was computed. Similarly, for the area under the precision-recall curve (**AUPRC**), the probabilities were used to construct the precision-recall curve, and the AUPRC value was calculated from the area under this curve.

PocketMiner encountered prediction errors for 22 structures from the test subset, leading to their exclusion from the evaluation. Additionally, PocketMiner cannot make predictions for multi-chain structures, which resulted in another 38 structures being removed from the test set. Thus, in total, 60 structures were excluded during PocketMiner evaluation, representing more than a quarter of the entire test set. Furthermore, during the evaluation of P2Rank, only single-chain structures were used. Consequently, three rounds of pLM-NN evaluation were conducted: the first using the entire test set to establish the benchmark^4^, the second for a direct comparison with PocketMiner, excluding the structures where PocketMiner was unable to make predictions, and the third for a direct comparison with P2Rank, excluding only structures with pockets spanning more than one chain.

The respective values are reported in Table 4. Remarkably, the fairly straightforward approach utilizing the train set from the CryptoBench dataset combined with the embeddings from the ESM2-3B model resulted in a neural network that matches or even surpasses the performance of the PocketMiner tool on the CryptoBench test set. Particularly, the neural network significantly outperforms PocketMiner in key metrics such as AUC and AUPRC, which are independent of the decision threshold selection.

**Table 4.** Performance of the benchmark method, PocketMiner, and P2Rank was evaluated across different subsets of the CryptoBench test set. In the first evaluation round, the benchmark method was assessed using the full CryptoBench test set (CB-full). In the second round, both the benchmark method and PocketMiner were evaluated on a subset that included only structures for which PocketMiner did not fail (CB-PM); see supplementary for details. In the third round, the benchmark method and P2rank were tested on a subset consisting solely of single-chain apo structures (CB-P2RANK-apo). Lastly, P2Rank was evaluated on holo structures (CB-P2RANK-holo), which are the counterparts of the apo structures from the CB-P2RANK-apo subset. The comparison between P2Rank’s performance on CB-P2RANK-apo and CB-P2RANK-holo highlights the performance drop when identifying cryptic binding sites using a method not specialized for detecting such sites.

| Method | Dataset | AUC | AUPRC | ACC | FPR | TPR | MCC | F1 Score |
| --- | --- | --- | --- | --- | --- | --- | --- | --- |
| pLM-NN | CB-full | 0.86 | 0.36 | 0.88 | 0.11 | 0.67 | 0.37 | 0.38 |
| pLM-NN | CB-PM | 0.88 | 0.43 | 0.88 | 0.11 | 0.71 | 0.41 | 0.42 |
| PocketMiner | CB-PM | 0.76 | 0.19 | 0.86 | 0.11 | 0.43 | 0.22 | 0.26 |
| pLM-NN | CB-P2RANK-apo | 0.88 | 0.42 | 0.89 | 0.10 | 0.70 | 0.40 | 0.41 |
| P2RANK | CB-P2RANK-apo | 0.81 | 0.21 | 0.85 | 0.14 | 0.62 | 0.27 | 0.28 |
| P2RANK | CB-P2RANK-holo | 0.89 | 0.34 | 0.85 | 0.15 | 0.84 | 0.38 | 0.35 |

Although PocketMiner achieves fairly competitive FPR values at the decision threshold set to 0.75, it also exhibits reduced TPR values. However, as shown in Figure 4, adjusting the decision thresholds to increase TPR also causes the FPR to rise steeply. This is particularly problematic for cryptic pocket predictions, given the imbalance between the number of binding and non-binding residues (as can be observed in Table 2 — on average, the binding residues only correspond to less than 5% of all residues in the whole protein). As a result, when selecting different decision thresholds, the high FPR combined with the imbalance between binding and non-binding residues could cause the number of false positives to significantly outnumber the true positives, which may reduce the usefulness of PocketMiner’s predictions.

**Fig. 4.**
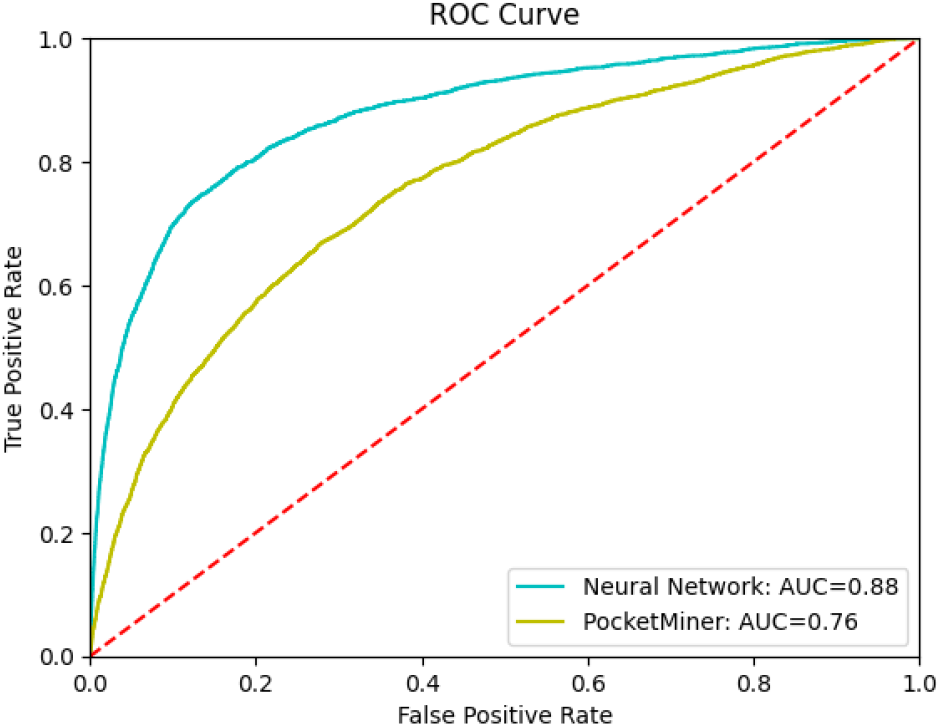
The performance of the benchmark method and PocketMiner on the test subset (CB-PM) is illustrated with the ROC curve. It needs to be emphasized that the ROC curve was constructed utilizing the probabilities generated by the neural network, as opposed to the statistics from Table 4, which were mostly computed using binary classified values of 0 and 1.

Although not specifically designed for CBS detection, P2Rank shows competitive performance on apo structures within the CryptoBench test set, a pattern consistent with findings from other studies [4, 5] on different CBS datasets. However, it does not surpass the benchmark method (pLM-NN), as shown in Table 4. As a validation step, P2Rank was also evaluated on holo structures^5^ to assess the performance difference of a non-CBS-specific method between apo and holo structures. In the case of structure-based methods, predicting cryptic binding sites is generally less challenging on holo structures, as their pockets are more clearly defined compared to their apo counterparts and align better with the holo-based training sets. Therefore, it is not surprising that P2Rank performs better on holo structures than on apo structures, a trend also observed in the aforementioned studies [4, 5] and also reflected in the results from the CryptoBench test set, as shown in Table 4 (comparing P2Rank performance on *CB-P2RANK-apo* vs. *CB-P2RANK-holo* dataset).

## Discussion

A universally accepted definition of what constitutes a CBS does not exist, as discussed in detail in Vajda et al. [1]. Typically, a pocket is considered cryptic if it binds a ligand in one form (holo) but not in another (apo). The RMSD-based crypticity criterion, as detailed in section Dataset Construction, aligns well with this definition since a large structural rearrangement (either closing or opening) is likely to prevent ligand binding. However, this criterion may fail to detect CBSs with small changes, such as a minor side chain rotation. While such modifications might prevent ligand binding, the overall site may not appear significantly altered. The previously introduced Cryptosite dataset has 35 structure pairs with a pocket RMSD smaller than 2 Å with the smallest difference as small as 0.19 Å. Figure 5 and Figure 6 give two examples from the original Cryptosite dataset. Figure 5 describes a structural pair where we do not see any substantial conformational change of the binding site, while Figure 6 shows an example of Cryptosite structure pair, where pocket RMSD is around 1 Å (similar to the previous example), but there is a substantial conformational change due to a change in conformation of one of the binding residues side chains that will likely affect the ligand binding. Although our definition of the cryptic binding sites might miss some genuine cryptic sites, the main focus of the benchmark is to minimize false positives (non-CBS regions labeled as CBS) rather than false negatives (CBS not labeled as CBS). Moreover, because our primary motivation for creating CryptoBench is to aid in the development of CBS prediction methods, a CBS definition that emphasizes large structural changes is appropriate. Detecting a binding site in an apo structure that significantly differs from the holo state is more challenging and thus aligns with our objectives. Finally, a crypticity criterion independent of the results of another prediction tool (such as a docking program) prevents the introduction of a bias or dependency on a certain set of parameters.

**Fig. 5.**
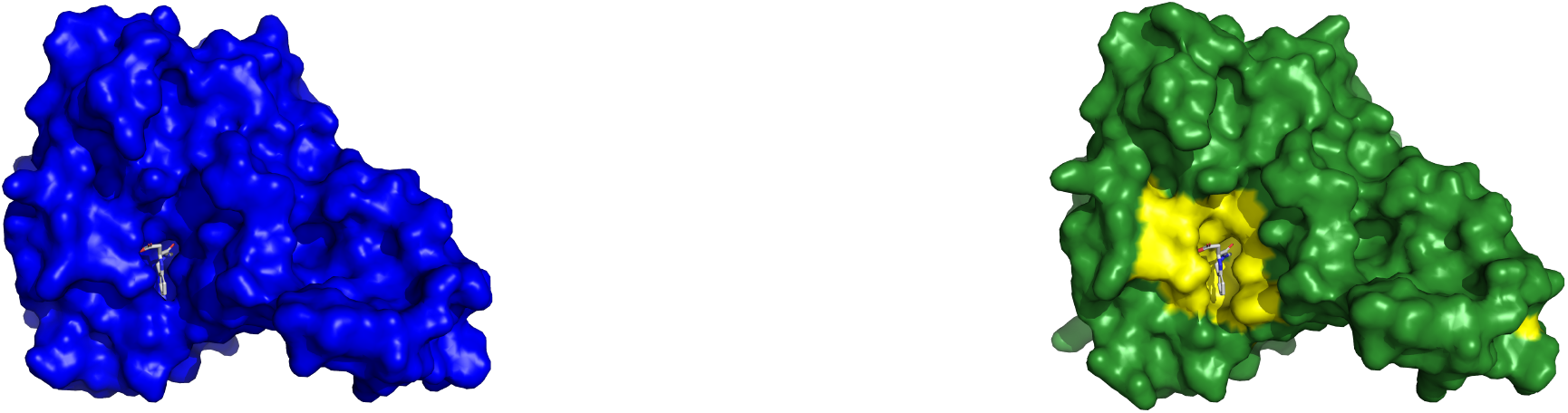
Example of a Cryptosite pair with pocket RMSD smaller than 2Å without a major change of the binding pocket. Bacterial esterase in the apo form (1qlw, left) and in the holo form (2wkw, right) with W22 ligand. Two structures were aligned and W22 from the holo structure is also shown in the aligned apo structure binding pocket to visualize the fit of the ligand into the apo structure. The binding site in both structures is almost identical.

**Fig. 6.**
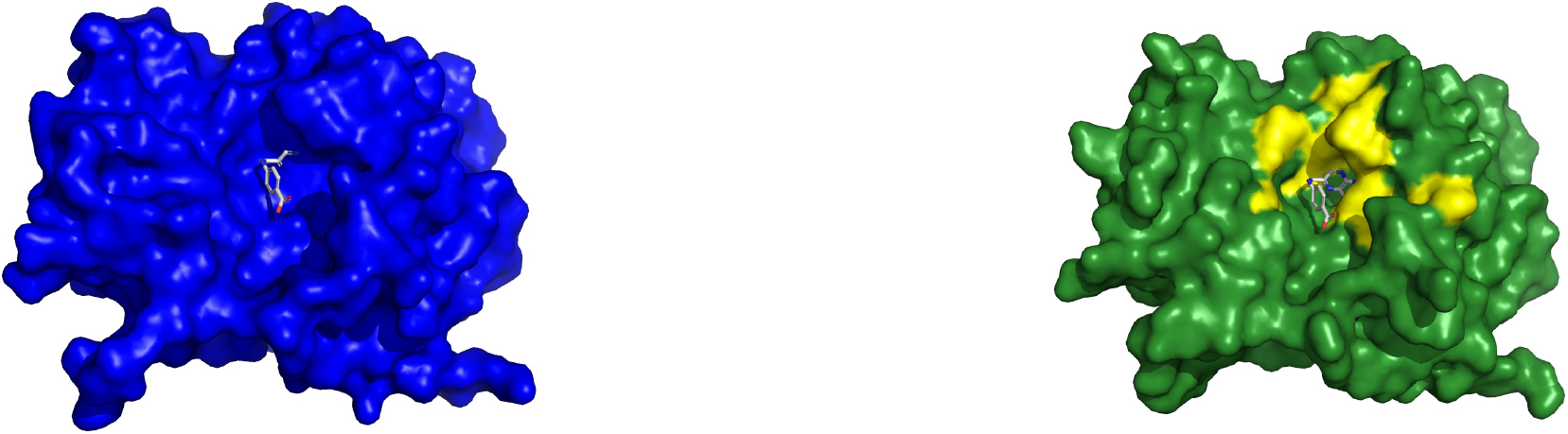
Example of Cryptosite pair with pocket RMSD smaller than 2Å with a major change of the binding pocket. Ricin in the apo form (1rtc, left) and in the holo form (1br6, right) with PT1 ligand. Two structures were aligned and PT1 from the holo structure is shown also in the aligned apo structure binding pocket to visualize the fit of the ligand into the apo structure. Tyr 80 sidechain conformation obstructs the ligand-binding site in the apo form.

To visually showcase the CryptoBench dataset, we give two examples with different RMSDs - one with RMSD close to the inclusion threshold and the other with a clear difference between the apo and holo states. Figure 7 shows cobyrinic acid a,c diamne synthatase with pocket RMSD 2.21 Å. The ANP ligand does not fit into the apo binding pocket due to a conformational change or residues 21-23. Figure 8 describes the difference of binding pockets of apo and holo structures of Cap-specific mRNA methyltransferase - the pocket RMSD of these structures is 3.03 Å. A loop consisting of binding site residues 277 to 280 has different conformation in apo structure resulting in a reduced size of the binding site. For aligned cartoon visualizations of the aforementioned apo-holo pairs, refer to the supplementary material, section 6.

**Fig. 7.**
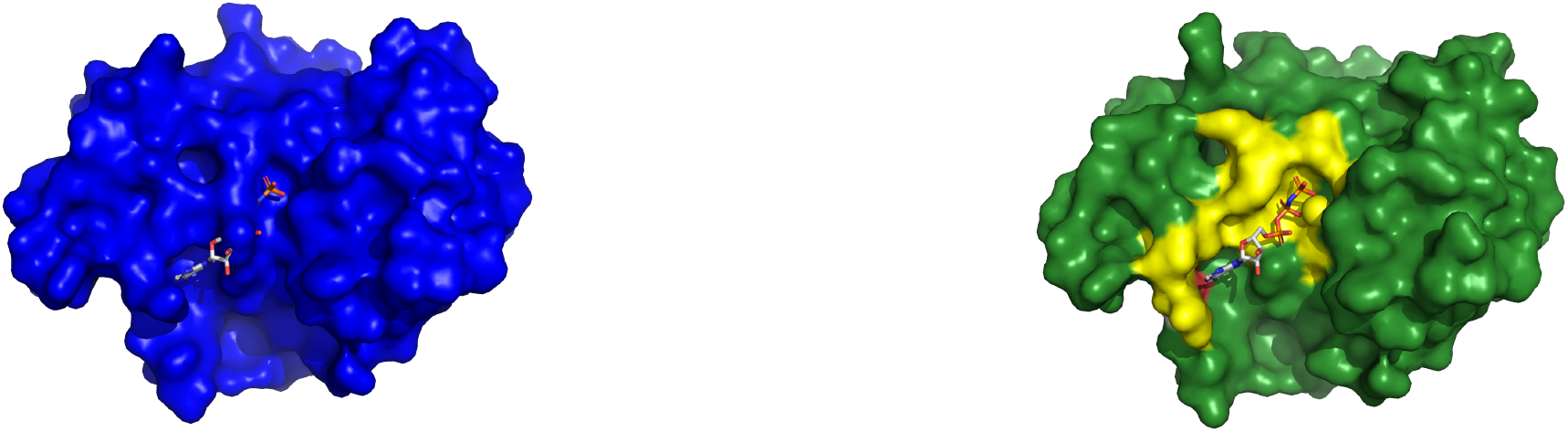
Example of a cryptic binding site from CryptoBench dataset. Cobyrinic acid a,c diamide synthase in the apo form (4pfs, left) and in the holo form (5if9, right) with ATP analog ANP. Two structures were aligned, and ANP from the holo structure is shown also in the aligned apo structure binding pocket to visualize the fit of the ligand into the apo structure. Residues Gly 21 to Ala 23 extend into one more turn of a helix in apo structure while they turn into a loop in the holo structure. This extra turn of helix in apo structure occupies the ligand-binding site.

**Fig. 8.**
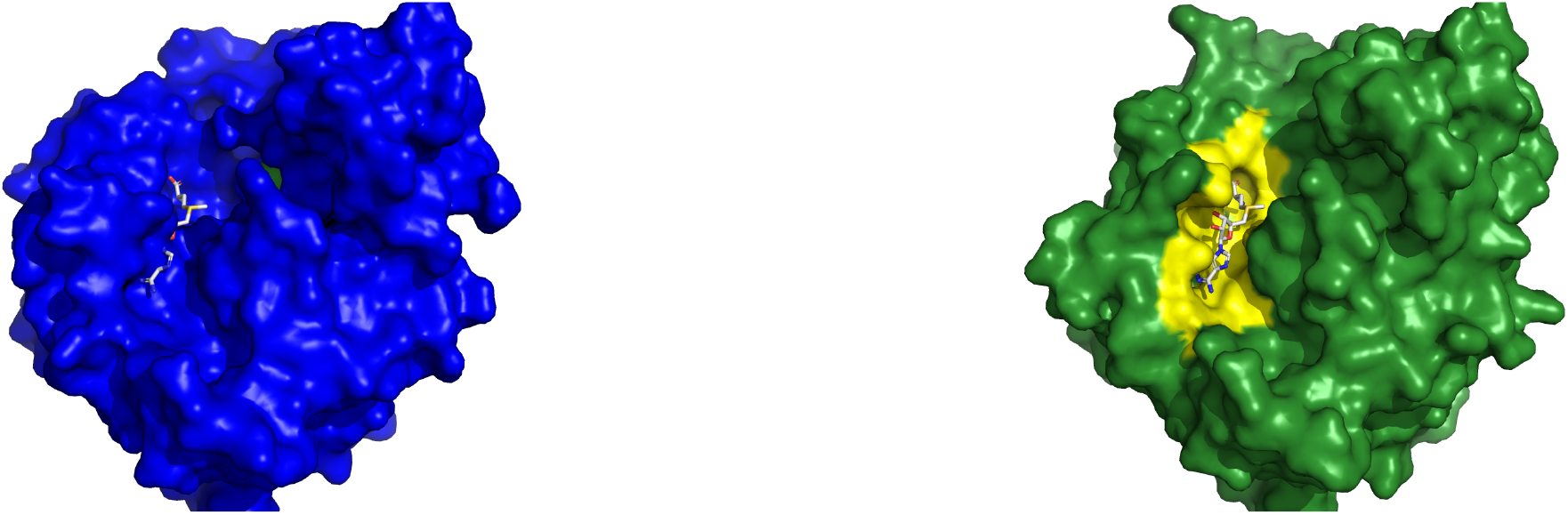
Example of a cryptic binding site from CryptoBench dataset. Cap-specific mRNA methyltransferase in the apo form (4n4a, left) and in the holo form (4n49, right) with SAM in the binding pocket. Two structures were aligned and SAM from the holo structure is shown also in the aligned apo structure binding pocket to visualize the fit of the ligand into the apo structure. The loop covering residues Ala 277 to Pro 280 has a different conformation in apo structure, which changes the binding pocket.

### Alternative Crypticity Metrics

During the creation of the dataset, other criteria were considered for inclusion in the filtering pipeline, in addition to those mentioned in the section Apo-holo pairs pool filtering. Records with a pocket RMSD larger than 6 Å were manually analyzed to determine if an upper bound on pocket RMSD was necessary. After visual inspection, it was concluded that even pockets with an RMSD greater than 6 Å are valid cryptic pockets; therefore, it was determined that an upper bound for pocket RMSD is unnecessary. Similarly, we considered setting an upper limit for the number of atoms in ligands to ensure the biological relevance of large ligands. We manually reviewed the list of apo-holo pairs where the number of heavy atoms in ligands exceeded 60. After visual inspection, we concluded that even large ligands remain biologically relevant, and therefore, we decided to keep them in the dataset.

Further, we have decided to keep even ligands that are covalently attached as they have proven to be relevant and successful in drug design [30].

## Conclusion

Leveraging the pocket RMSD metric, we curated the CryptoBench dataset, filtering apo-holo pairs and identifying cryptic pockets in PDB. The dataset’s structural diversity enables fair validation of machine learning models for cryptic binding site prediction. We also provided an evaluation benchmark comprising representatives of sequence- and structure-based prediction methods. We believe that CryptoBench will prove valuable to the scientific community, further catalyzing the development of more accurate and reliable methods for identifying both general and cryptic binding sites.

## Availability

The CryptoBench dataset is available on Open Science Framework - https://osf.io/pz4a9/ [31].

## Competing interests

No competing interest is declared.

## Author contributions statement

D.H. conceived the research, V.S. and D.H. conceived the experiments, V.S. created the benchmark and conducted the experiments, C.F. provided the apo-holo data and together with M.N consulted the data and biology aspects of the research, R.K. provided the P2Rank evaluation, V.S. and D.H. wrote the manuscript, all authors discussed the results and reviewed the manuscript.

## Acknowledgments

This work was supported by the Czech Science Foundation (GAC?R) grant number 23-07349S, and the ELIXIR CZ Research Infrastructure (ID LM2018131, MEYS CR).

Computational resources were provided by the e-INFRA CZ project (ID:90254), supported by the Ministry of Education, Youth and Sports of the Czech Republic.

## Footnotes

1 A subset of AHoJ-DB was used — records with low resolution (*>* 2.5 Å) or with pockets containing unobserved residues were filtered out. That is, the subset consisted of 4,683,968 apo-holo pairs.

2 The original P2Rank ignored group list also included sugars MAN, GLC, and NAG. However, these sugars are biologically relevant, as evidenced by cases like the 1esw-5jiw apo-holo pair involved in starch synthesis. Additionally, other studies like AlphaFold3 [23] chose to include these sugars.

3 For multiple-chain records, each chain was considered separately to calculate the average number of observed residues.

4 In the first round of the evaluation, for structures with multiple chains, each chain was treated as an individual entity during the process.

5 Any apo structure in CryptoBench can be associated with more than one holo structure. In the case of evaluating P2Rank on holo structures, a single holo structure needed to be selected for a fair comparison. Therefore, holo structure with the largest pocket RMSD was selected; see supplementary, section 7.

## Notes

https://osf.io/pz4a9/

### Competing Interest Statement

The authors have declared no competing interest.

