## Supplementary Document for "CryptoBench: Cryptic protein-ligand binding sites dataset and benchmark"

August 2024

#### 1 Surface metrics

The primary crypticity metrics we considered included pocket solvent accessible surface area (SASA), pocket molecular surface, pocket volume, and all-atom pocket RMSD. Figure 1 illustrates the difference between SASA and molecular surface.

#### 2 Missing structures in the evaluation

As stated in the manuscript, PocketMiner encountered prediction errors for certain structures from the CryptoBench test dataset. Furthermore, structures containing multiple chains had to be excluded, as PocketMiner cannot operate with multi-chain structures. The list of all the excluded structures is provided here:

- No prediction due to prediction error: 5ighA, 1se8A, 5yj2C, 3bjpA, 4fkmB, 2phzA, 3mwgB, 3t8bA, 4dncB, 2dfpA, 7nbcCCC, 5acvB, 4bg8A, 1h13A, 4jaxF, 2vqzF, 3uyiA, 2vyrA, 1x2gC, 7c48A, 3pfpA, 7qzrD, 1k47D
- No prediction due to multi-chain structure: 4x19 M-O-P, 1fd4 G-H, 5aon A-B, 1fe6 A-D, 7nbc AAA-CCC, 5wm9 B-C, 3la7 A-B, 3bk9 E-H,

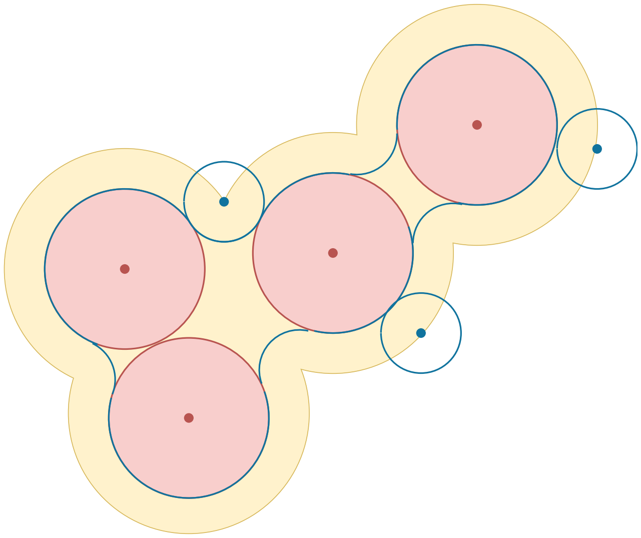

Figure 1: Illustration of solvent accessible surface area (yellow curve), molecular surface (blue curve). Both areas are formed by a spherical probe rolling over the protein's surface. The red circles represent van der Waals surfaces of the atoms, and the blue circles depict the spherical probe. Throughout our study, the value of 1.4 Å is used as the probe radius.

2czd A-B, 3ve9 A-B, 5ujp A-B, 7np0 A-B, 1tmi A-B, 3hrm A-B, 2nt1 A-B, 2xdo A-C, 3b1o A-B, 5h8k I-J, 3kjr A-B, 8gxj B-C, 2idj A-B, 5m7r A-B, 4z0y B-D-F-H, 2zcg A-B, 5n49 A-B, 8hc1 d-h, 3lnz C-O, 1xxo A-B, 8h49 A-B, 6syh A-B, 1g1m A-B, 3pfp A-B, 5dy9 H-I, 4p32 A-B, 2huw A-B, 5gmc A-B, 1r3m A-B, 3x0x E-G.

Similarly, P2Rank was evaluated on the single-chain structures only. The list of excluded structures for P2Rank evaluation is provided here:

- 4x19 M-O-P, 1fd4 G-H, 5aon A-B, 1fe6 A-D, 7nbc AAA-CCC, 5wm9 B-C, 3la7 A-B, 3bk9 E-H, 2czd A-B, 3ve9 A-B, 5ujp A-B, 7np0 A-B, 1tmi A-B, 3hrm A-B, 2nt1 A-B, 2xdo A-C, 3b1o A-B, 5h8k I-J, 3kjr A-B, 8gxj B-C, 2idj A-B, 5m7r A-B, 4z0y B-D-F-H, 2zcg A-B, 5n49 A-B, 8hc1 d-h, 3lnz C-O, 1xxo A-B, 8h49 A-B, 6syh A-B, 1g1m A-B, 3pfp A-B, 5dy9 H-I, 4p32 A-B, 2huw A-B, 5gmc A-B, 1r3m A-B, 3x0x E-G.

The list is identical with the list of excluded multi-chain structures in PocketMiner.

### 3 The mapping from the PDB-based annotations to the Uniprot-based embeddings

As mentioned in the manuscript, the embeddings were initially generated from full UniProt sequences. Therefore, only those embeddings corresponding to observed residues in the PDB structures were used. The mapping from UniProt sequences to PDB observed residues was facilitated through the PDB's API[1]. Consequently, minor inconsistencies may exist in the final statistics; for instance, slight variations in protein length between PocketMiner data and NN data. Nonetheless, these differences are negligible; for instance, the average sequence length for pLM-NN evaluated on test subset consisting only of structures not failing on PocketMiner (CB-PM) is 299, while for PocketMiner, it's 302. This disparity arises from some PDB residues being absent in the UniProt records.

### 4 Additional graphs

The graphs presented in the manuscript depict both the decrease and increase of the molecular surface, SASA, and volume metrics between the apo-holo state. However, for readers interested in observing the violin plots illustrating the absolute values, please refer to Figure 2. In this case, the formula used to calculate the difference is:

$$difference = \frac{|apo\_value - holo\_value|}{pocket\_length} \tag{1}$$

### 5 Low-RMSD cryptic pockets from PocketMiner

To enhance our confidence in CryptoBench's 2Å RMSD threshold for determining crypticity, we measured how many of the PocketMiner dataset cryptic pockets match the criterion. We found that 6 out of 38 pairs from the PocketMiner dataset do not meet the threshold. Within this subset, 2 pairs (apo:

[1]For more details, see `https://www.ebi.ac.uk/pdbe/graph-api/pdbe_doc/#api-Residue-GetPDBResidueRangeAnnotations`

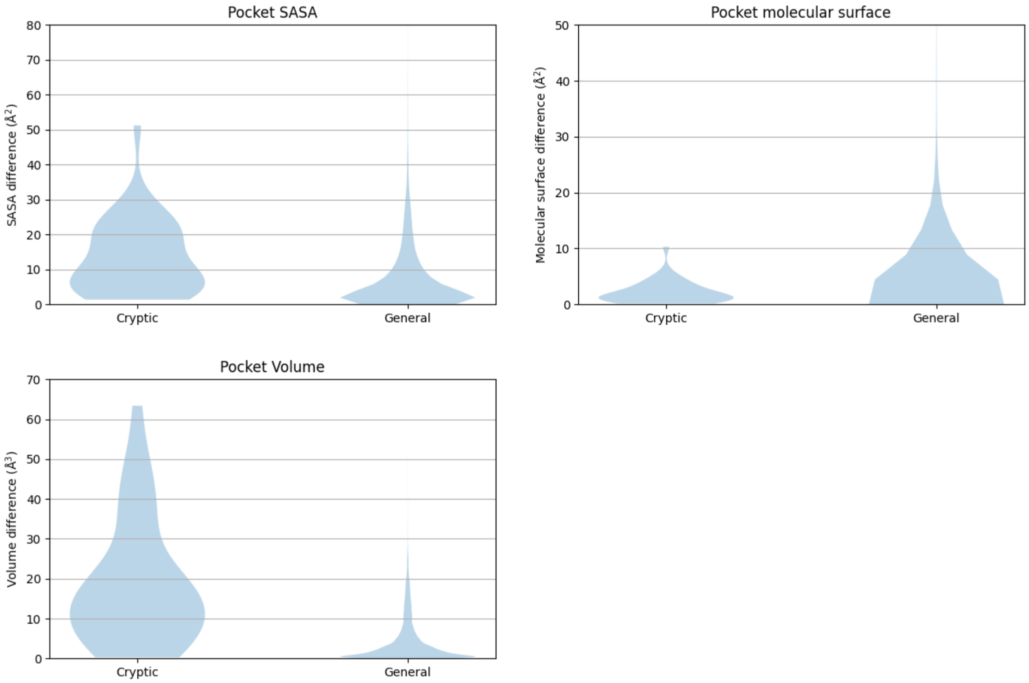


Figure 2: Violin plots illustrating the absolute difference of the metrics between the apo and holo state.

*5h9a, 1s2o*) miss the threshold only by a narrow margin, with pocket RMSD values exceeding 1.8 Å, two pairs (apo: *4w51, 6e5d*) had pocket RMSD values falling within the interval of $[1.5, 1.8]$ Å and only 2 pairs exhibited pocket RMSD values lower than 1.5 Å (apo: *4i92* and *3ppn*).

The main text of the manuscript contains discussion and visualization of the *4i92* apo structure. Here, we also provide examples of *4w58* (Figure 3) and *5h9a* (Figure 4).

### 6 Cartoon visualizations

For each apo-holo pair discussed in the Discussion section, an aligned and cartoon-visualized version was prepared. In Cryptosite, the *1qlw-2wkw* pair is shown in Figure 5, and the *1rtc-1br6* pair in Figure 6. For CryptoBench, the *4pfs-5if9* pair is shown in Figure 7, and the *4n4a-4n49* pair in Figure 8.

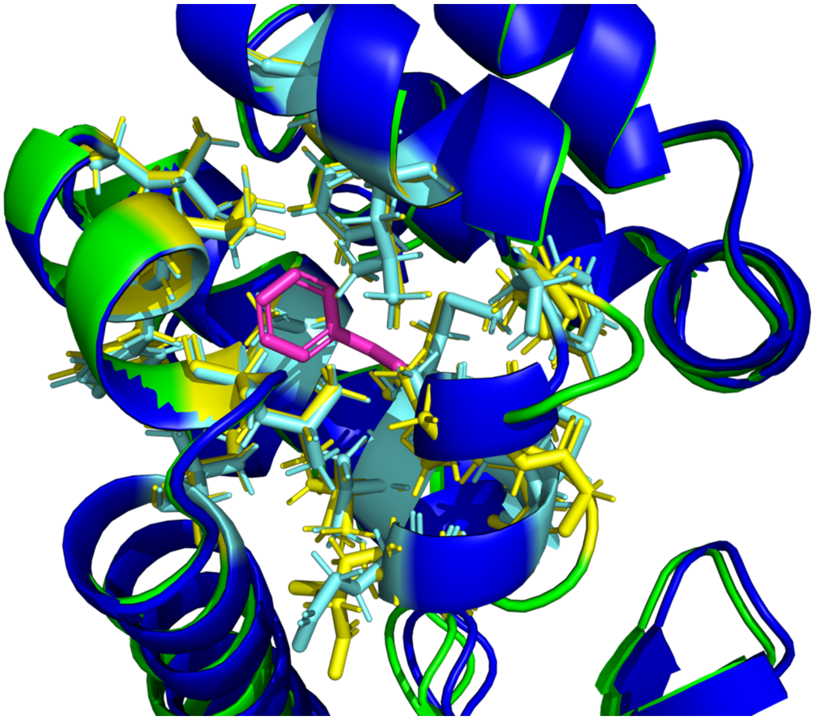

Figure 3: A superposition of an apo and a holo structure of T4 lysozyme with limited conformational change. The figure depicts a cryptic binding site as described in PocketMiner: apo structure (4w51) – blue structure with light blue sticks describing the binding site residues as identified in the holo structure (4w58) - green structure with yellow sticks describing binding sites of pentylbenzene (in magenta). The binding sites overlap very closely – only Glu 108 in the bottom left corner diverges slightly.

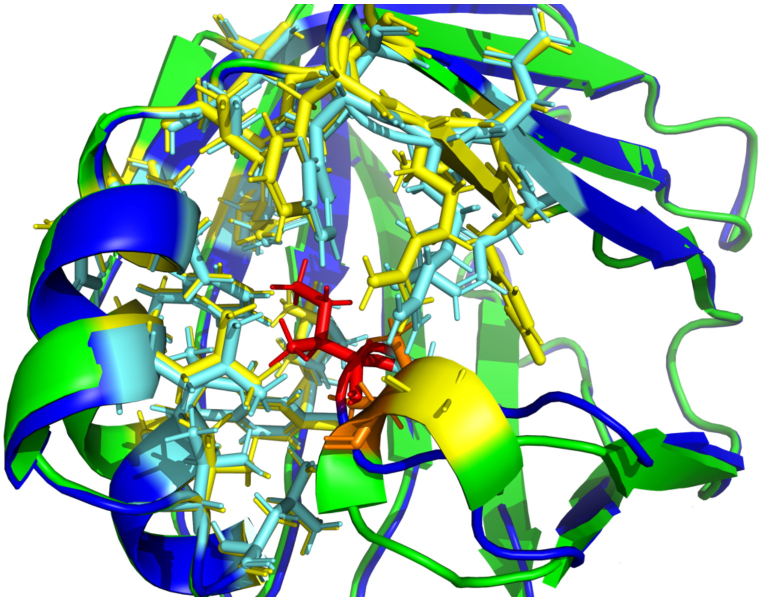

Figure 4: A superposition of an apo and a holo structure of retinol-binding protein 1 with significant conformational change. A figure depicts a cryptic binding site as described in PocketMiner: apo structure (5h9a) – blue structure with light blue sticks describing the binding site residues as identified in the holo structure (6e5l) - green structure with yellow sticks describing binding sites of HVD (not shown for clarity). Residue Ile 77 (red in apo and orange in holo structure) undergoes a substantial conformational change. It is moved out from the binding pocket in holo structure, but in the apo structure, the side chain of isoleucine interferes with the binding pocket of the ligand.

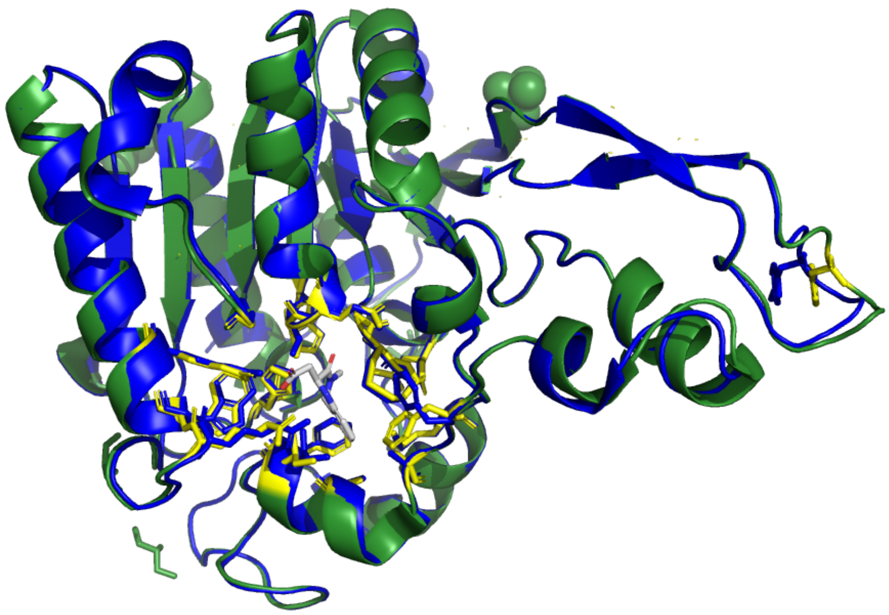

Figure 5: Example of Cryptosite pair with pocket RMSD smaller than 2Å without a a major change of the binding pocket. Structure superposition of esterase in the apo form (1qlw, blue) and the holo form (2wkw, green) with W22 ligand. The binding site is in yellow. The ligand is gray. Figure was prepared in PyMOL.

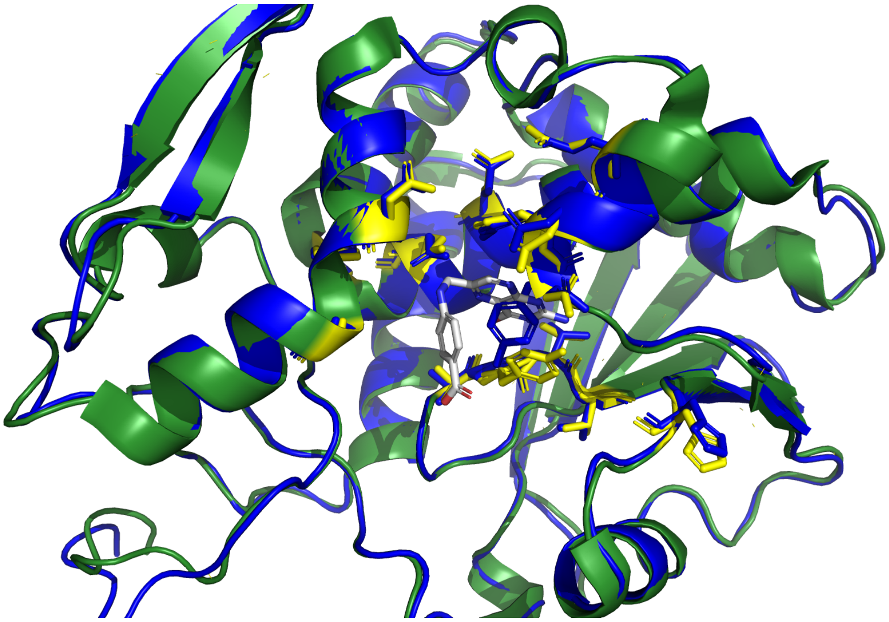

Figure 6: Example of Cryptosite pair with pocket RMSD smaller than 2Å with a major change of the binding pocket. Structure superposition of ricin in the apo form (1rtc, blue) and in the holo form (1br6, green) with PT1 ligand. The binding site is in yellow. The ligand is gray. Figure was prepared in PyMOL.

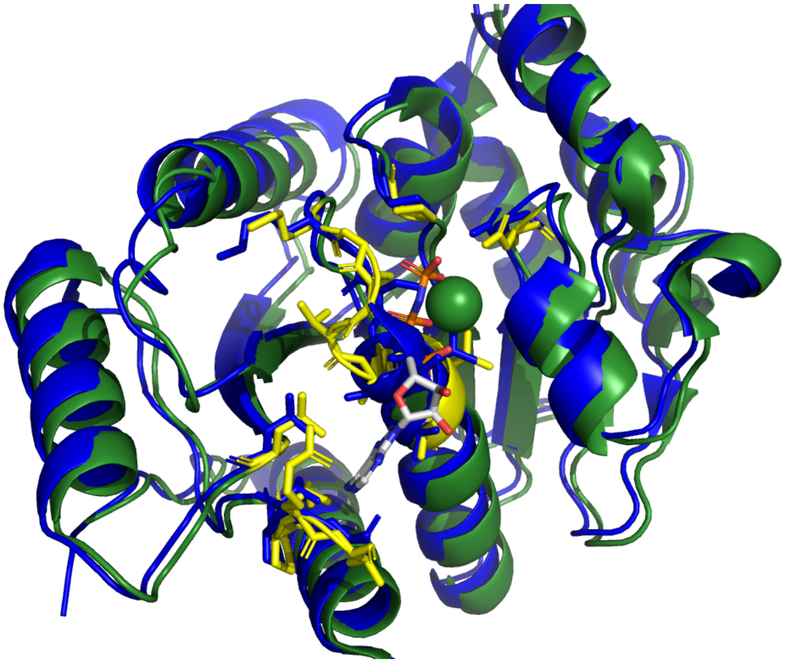

Figure 7: Structure superposition of cobyrinic acid a,c diamide synthase in the apo form ( 4pfs, blue) and in the holo form (5if9, green) with ATP analog ANP. The binding site is in yellow. The ligand is gray. Figure was prepared in PyMOL.

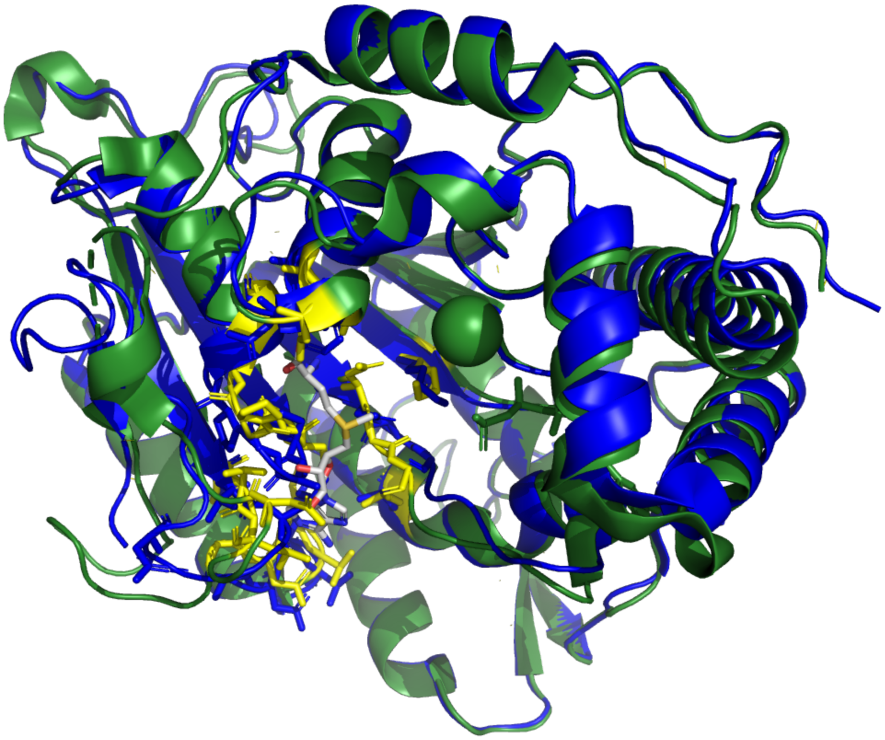

Figure 8: Structure superposition of Cap-specific mRNA methyltransferase in the apo form (4n4a, blue) and in the holo form (4n49, green) with SAM in the binding pocket. The binding site is in yellow. The ligand is gray. Figure was prepared in PyMOL.

### 7 HOLO structures for P2Rank evaluation

The HOLO structures, which were used for P2Rank evaluation, are flagged in the dataset using `is_main_holo_structure` attribute.
